# Addressing the looming identity crisis in single cell RNA-seq

**DOI:** 10.1101/150524

**Authors:** Megan Crow, Anirban Paul, Sara Ballouz, Z. Josh Huang, Jesse Gillis

**Affiliations:** Cold Spring Harbor Laboratory, One Bungtown Road, Cold Spring Harbor, NY 11724, USA

**Keywords:** single cell RNA-sequencing, neural diversity, transcriptome, interneuron, cell type, replicability, bioinformatics

## Abstract

Single cell RNA-sequencing technology (scRNA-seq) provides a new avenue to discover and characterize cell types, but the experiment-specific technical biases and analytic variability inherent to current pipelines may undermine the replicability of these studies. Meta-analysis of rapidly accumulating data is further hampered by the use of *ad hoc* naming conventions. Here we demonstrate our replication framework, MetaNeighbor, that allows researchers to quantify the degree to which cell types replicate across datasets, and to rapidly identify clusters with high similarity for further testing. We first measure the replicability of neuronal identity by comparing more than 13 thousand individual scRNA-seq transcriptomes, sampling with high specificity from within the data to define a range of robust practices. We then assess cross-dataset evidence for novel cortical interneuron subtypes identified by scRNA-seq and find that 24/45 cortical interneuron subtypes have evidence of replication in at least one other study. Identifying these putative replicates allows us to re-analyze the data for differential expression and provide lists of robust candidate marker genes. Across tasks we find that large sets of variably expressed genes can identify replicable cell types and subtypes with high accuracy, suggesting a general route forward for large-scale evaluation of scRNA-seq data.

---

Single cell RNA-sequencing (scRNA-seq) has emerged as an important new technology enabling the dissection of heterogeneous biological systems into ever more refined cellular components. One popular application of the technology has been to try to define novel cell subtypes within a given tissue or within an already refined cell class, as in the lung^1^, pancreas^2-5^, retina^6, 7^, or others^8-10^. Because they aim to discover completely new cell subtypes, the majority of this work relies on unsupervised clustering, with most studies using customized pipelines with many unconstrained parameters, particularly in their inclusion criteria and statistical models^7, 8, 11, 12^. While there has been steady refinement of these techniques as the field has come to appreciate the biases inherent to current scRNA-seq methods, including prominent batch effects^13^, expression drop-outs^14, 15^, and the complexities of normalization given differences in cell size or cell state^16, 17^, the question remains: how well do novel transcriptomic cell subtypes replicate across studies?

In order to answer this, we turned to the issue of cell diversity in the brain, a prime target of scRNA-seq as neuron diversity is critical for construction of the intricate circuits underlying brain function. The heterogeneity of brain tissue makes it particularly important that results be assessed for replicability, while its popularity as a target of study makes this goal particularly feasible. Because a primary aim of neuroscience has been to derive a taxonomy of cell types^18^, already more than twenty single cell RNA-seq experiments have been performed using mouse nervous tissue^19^. Remarkable strides have been made to address fundamental questions about the diversity of cells in the nervous system, including efforts to describe the cellular composition of the cortex and hippocampus^11, 20^, to exhaustively discover the subtypes of bipolar neurons in the retina^6^, and to characterize similarities between human and mouse midbrain development^21^. This wealth of data has inspired attempts to compare data^6, 12, 20^ and more generally in the single cell field there has been a growing interest in using batch correction and related approaches to fuse data across replicate samples or across experiments^6, 22, 23^. Historically, data fusion and modeling of experimental confounds have been necessary steps precisely where individual experiments are underpowered or results do not replicate without correction^24-26^ but even sophisticated approaches to merge data come with their own perils^27^. The technical biases of scRNA-seq have motivated interest in correcting them as a seemingly necessary fix, yet evaluation of whether results replicate in the first place remains largely unexamined and no systematic or formal method has been developed for accomplishing this task.

To address this gap in the field, we propose a simple, supervised framework, MetaNeighbor (**meta**-analysis via **neighbor** voting), to assess how well cell type-specific transcriptional profiles replicate across datasets. Our basic rationale is that if a cell type has a biological identity rooted in the transcriptome then knowing its expression features in one dataset will allow us to find cells of the same type in another dataset. We make use of the cell type labels supplied by data providers, and assess the correspondence of cell types across datasets by taking the following approach (see schematic, Figure 1):

1) We calculate correlations between all pairs of cells that we aim to compare across datasets based on the expression of a set of genes. This generates a network where each cell is a node and the edges are the strength of the correlations between them.
2) Next, we do cross-dataset validation: we hide all cell type labels (‘identity’) for one dataset at a time. This dataset will be used as our test set. Cells from all other datasets remain labeled, and are used as the training set.
3) Finally, we predict the cell type labels of the test set: we use a neighbor voting algorithm to predict the identity of the held-out cells based on their similarity to the training data.

**Figure 1.**
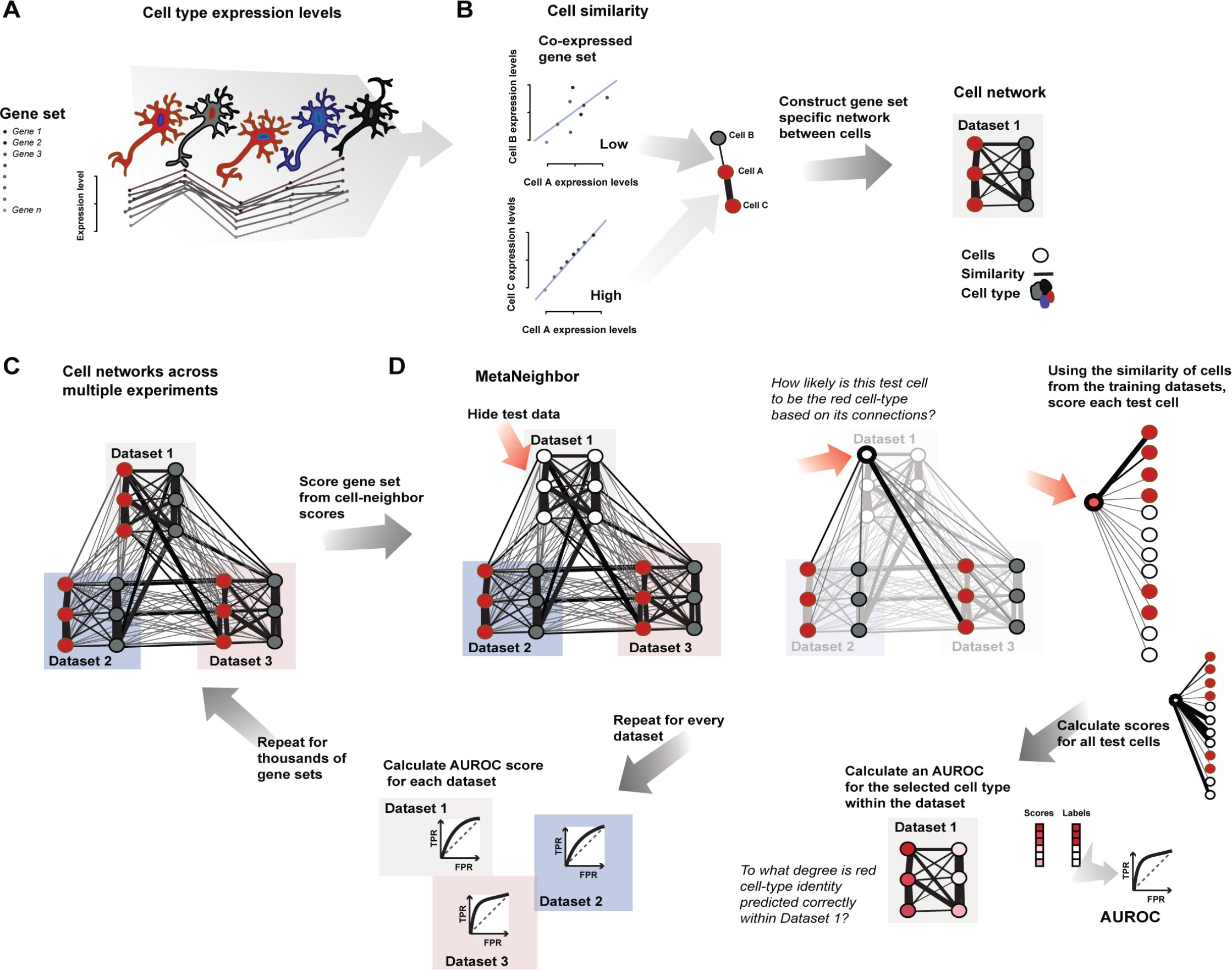
MetaNeighbor quantifies cell type identity across experiments. **A** – Schematic representation of gene set co-expression across individual cells. Cell types are indicated by their color. **B** – Similarity between cells is measured by taking the correlation of gene set expression between individual cells. On the top left of the panel, gene set expression between two cells, A and B, is plotted. There is a weak correlation between these cells. On the bottom left of the panel we see the correlation between cells A and C, which are strongly correlated. By taking the correlations between all pairs of cells we can build a cell network (right), where every node is a cell and the edges represent how similar each cell is to each other cell. **C** – The cell network that was generated in B can be extended to include data from multiple experiments (multiple datasets). The generation of this multi-dataset network is the first step of MetaNeighbor. **D** – The cross-validation and scoring scheme of MetaNeighbor is demonstrated in this panel. To assess cell type identity across experiments we use neighbor voting in cross-validation, systematically hiding the labels from one dataset at a time for testing. Cells within the test set are predicted as similar to the cell types from other training sets using a neighbor voting formalism. Whether these scores prioritize cells as the correct type within the dataset determines the performance, expressed as the AUROC. In other words, comparative assessment of cells occurs only within a dataset, but is based only on training information from outside that dataset. This is then repeated for all gene sets of interest.

Conceptually, this resembles approaches for the validation of sample clustering^28, 29^, which have primarily been applied to compare microarray results with respect to tumor subtyping^30, 31^. Our method builds on these ideas, adapting and applying them for the first time to the question of cell identity in single cell RNA-seq, and specifically exploiting the patterns of co-expression believed to drive results^32^. Because our implementation is extremely fast, this approach readily permits carefully defined control experiments to investigate the data features that drive high performance, such as the dependence on expression variability, gene set size, rarity of cell types or subtlety of transcriptional identity.

We evaluate the replicability of cell type transcriptional identity by taking sequential steps according to the basic taxonomy of brain cells: first classifying neurons vs. non-neuronal cells across eight single cell RNA-seq studies, then classifying cortical inhibitory neurons vs. excitatory neurons, and for our final step, we align interneuron subtypes across three studies. With detailed control experiments and empirical modeling, we validate the use of highly variable genes for cross-dataset cell identification, a common approach for feature selection within individual experiments^4, 33-35^. Testing hundreds of gene sets, we find strong replication of neuronal identity when compared to non-neurons, and excitatory vs. inhibitory neurons, even across widely varying techniques such as nuclear RNA-sequencing or Drop-seq. Furthermore, we find that cortical interneuron subtypes show clear lineage-specific structure, and we readily identify 11 subtypes that appear to replicate across datasets, including Chandelier cells and five novel subtypes defined by transcriptional clustering in previous work. Meta-analysis of differential expression across these highly replicable cortical interneuron subtypes correctly identified canonical marker genes such as parvalbumin and somatostatin, as well as new candidates which may be used for improved molecular genetic targeting, and to understand the diverse phenotypes and functions of these cells.

## Results

### Assessing neuronal identity with MetaNeighbor

We aimed to measure the replicability of cell identity across tasks of varying specificity. Broadly, these are divided into tasks where we are recapitulating known cell identities, and ones where we are measuring the replicability of novel cell identities discovered in recent research. The former class of task is the focus of this subsection: first, by assessing how well we distinguish neurons from non-neuronal cells (“task one”), and next assessing the discriminability of excitatory and inhibitory neurons (“task two”). As detailed in the methods, MetaNeighbor outputs a performance score for each gene set and task. This score is the mean area under the receiver operator characteristic curve (AUROC) across all folds of cross-dataset validation, and it can be interpreted as the probability that we will rank a positive higher than a negative. For example, if given only information from other (training) datasets labeling neurons and non-neurons, and asking the algorithm to identify neurons within a given (testing) dataset, the AUROC is the probability a neuron will be ranked above a non-neuron. Importantly, there is no labeling within the dataset being assessed; only signals which are true from one dataset to the next can contribute to performance. The AUROC varies between 0 and 1, with 1 being perfect classification, 0.5 meaning that we have performed as well as if we had randomly guessed the cell’s identity (null), and 0.9 or above being extremely high. Low scores (0-0.3) can be interpreted with as much confidence as high scores, and mean that, for example, a neuron is definitely not a non-neuron. Comparison of scores across gene sets allows us to discover their relative capacity to discriminate cell types.

As described above, in task one we assessed how well we could identify neurons and non-neuronal cells across eight datasets with a total of 13928 cells (Supplementary Table 1). Although this was designed to be fairly simple, we were interested to discover that AUROC scores were significantly higher than chance for all gene sets tested, including all randomly chosen sets (AUROC_all sets_=0.80 ± 0.1, Figure 2A). A bootstrapped sampling of the datasets showed a trend toward increased performance with the inclusion of additional training data, indicating that we are recognizing an aggregate signal across datasets (Supplementary Figure 1). However, the significant improvement of random sets over the null (i.e., AUROC=0.5) means that prior knowledge about gene function is not required to differentiate between these cell classes. Randomly chosen sets of genes have decidedly non-random expression patterns that enable discrimination between cell types. This is particularly surprising in the context of cross-dataset assessment, where the low-dimensionality of cell identity observed within laboratories^36^ is confounded by the even lower-dimensionality of experimental identity, even if controlled by within-lab ranking. This result recalls the startling finding by Venet *et al*. that “Most random gene expression signatures are related to breast cancer outcome”^37^; cell identity appears to be as clearly ascertainable.

**Figure 2.**
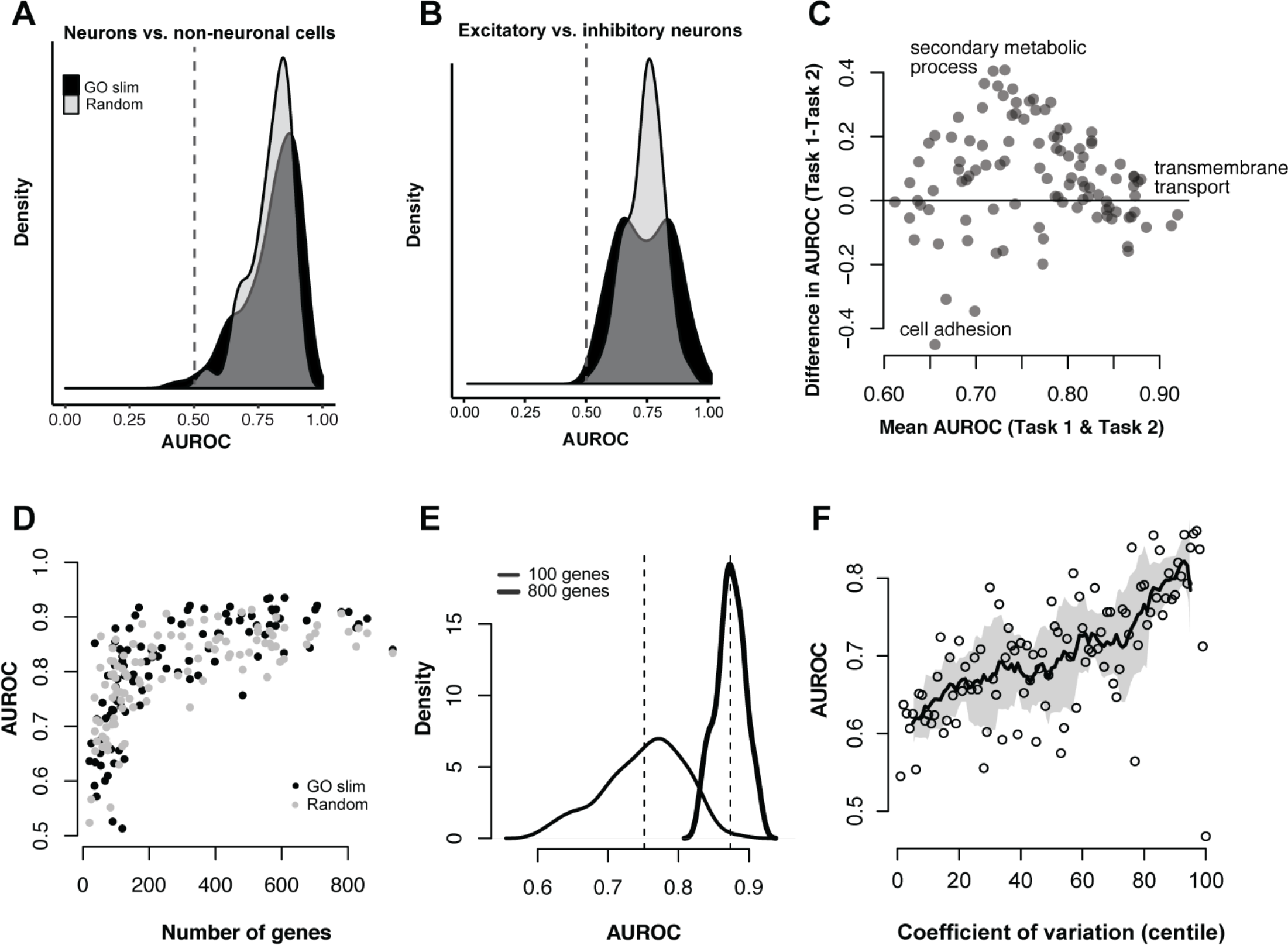
Cell type identity is widely represented in the transcriptome. **A & B** – Distribution of AUROC scores from MetaNeighbor for discriminating neurons from non-neuronal cells (“task one”, A) and for distinguishing excitatory vs. inhibitory neurons (“task two”, B). GO scores are in black and random gene set scores are plotted in gray. Dashed grey lines indicate the null expectation for correctly guessing cell identity (AUROC=0.5). For both tasks, almost any gene set can be used to improve performance above the null, suggesting widespread encoding of cell identity across the transcriptome. **C** – Comparison of GO group scores across tasks. GO groups at the extremes of the distribution are labeled. Most gene sets have higher performance for Task one, and a number of groups have high performance for both tasks (e.g., transmembrane transport). **D** – Task one AUROC scores for each gene set are plotted with respect to the number of genes. A strong, positive relationship is observed between gene set size and AUROC score, regardless of whether genes were chosen randomly or based on shared functions. **E** – Distribution of AUROC scores for task one using 100 sets of 100 randomly chosen genes, or 800 randomly chosen genes. The mean AUROC score is significantly improved with the use of larger gene sets (mean 100 = 0.80 +/− 0.05, mean 800 = 0.90 +/− 0.03). **F** – Relationship between AUROC score and coefficient of variation. Task one was re-run using sets of genes chosen based on mean coefficient of variation across datasets. A strong positive relationship was observed between this factor and performance (r_s_ ∼0.67).

Task two aimed to assess how well we could discriminate between cortical excitatory and inhibitory neurons across four studies with a total of 2809 excitatory and 1162 inhibitory neurons^11, 12, 20, 38^. Similar to our previous results, we saw that AUROC scores were significantly higher than chance (AUROC=0.69 ± 0.1, Figure 2B). While performance is higher than chance for both tasks, it is unclear whether the same gene sets are useful for distinguishing between neurons and non-neurons and between excitatory and inhibitory neurons. Comparing GO group performance across these two tasks we find that a handful of gene sets have high performance for both tasks (e.g., GO:0055085 transmembrane transport, AUROC>0.85, Figure 2C), while many GO groups show divergent performance. For example, we find that GO:0019748 (secondary metabolic process) is only useful for distinguishing between neurons and non-neurons, but not at all for distinguishing between the two neuron classes (AUROC_Task1_=0.94 vs. AUROC_Task2_=0.53), perhaps due to cell cycling among non-neuronal cells. On the other extreme, we find that GO:0040011 (cell adhesion) is only useful for distinguishing between neuron classes but not between neurons and non-neuronal cells (AUROC_Task1_=0.43 vs. AUROC_Task2_=0.88), which is in line with previous work that has found that cell adhesion factors show neuron-type specific expression^39, 40^. These results indicate some degree of functional specificity for gene set performance, but the near equivalent performance of randomly chosen gene sets suggests that transcriptional differences are likely to be encoded in a large number of genes, in line with previous observations^41^. The properties of high performing sets are investigated in the following section.

### Characterizing features associated with high performance

Consistent with the view that a large fraction of transcripts are useful for determining cell identity, we found a positive dependency of AUROC scores on gene set size, regardless of whether genes within the sets were randomly selected or shared some biological function (Figure 2D). This was further supported by a comparison of scores for task one when using randomly chosen sets of genes constrained to a given size. Here we used set sizes of 100 or 800, similar to the extremes of the distribution of set sizes used in the GO analysis. AUROC score distributions and means were significantly different between gene sets of different sizes, with sets of 100 genes having lower scores but higher variability in performance, whereas sets of 800 genes are more restricted in variance and give higher performance on average (Figure 2E, AUROC_100_=0.75 ± 0.06, AUROC_800_=0.87 ± 0.02, p<2.2E-16, Wilcoxon rank sum test). The variability in performance observed while keeping set size constant suggests that even in random sets, there are transcriptional features that contribute to cell identity. We delved into this further by comparing AUROC scores across gene sets chosen based on coefficient of variation, as MetaNeighbor relies on co-variation between genes to detect differences in cell type profiles. We performed task one again using these gene sets and found a strong positive relationship between variance and our ability to classify cells (Figure 2F, r_s_=0.67), though interestingly, genes in the top centile were completely uninformative (AUROC=0.47). Taken together, these observations support the idea that transcriptional identity is broadly encoded across many genes, and suggests that it should be straightforward to select an informative gene set that takes advantage of properties associated with high performance. Testing our capacity to detect and exploit this signal requires us to refine the cell classes that we are characterizing, ideally beyond what is present in existing data to anticipate a wide range of use cases.

### Empirical modeling to determine precision

Our ultimate aim is to identify all replicable cell types across datasets, some of which may be rare and/or only subtly different from other cell types. To assess the ability of MetaNeighbor to identify cell types in these more realistic scenarios, we set up an empirical model for cell type rarity and subtlety (schematic Figure 3A), using inhibitory and excitatory neuron datasets with >100 cells for each type as these allow us to model cell type incidence down to 1%^11, 12, 20^. To address the impact of rarity on MetaNeighbor’s performance, we alter the incidence of excitatory neurons to be within our observed range of subtype incidences, repeatedly sampling different combinations of cells to obtain mean performance estimates. Transcriptional subtlety is captured by only permitting a fraction of transcripts to vary between the two cell types. This treats transcriptional subtlety almost identically to a rare cell type, but in the dimension of transcripts rather than cells: a rare cell type is one in which only a few differing cells are present and a subtle transcriptional identity is one in which only a few differing genes are present. Subtlety is modeled by swapping out, e.g., the same 95% of the transcriptional profiles across all excitatory cell transcriptional data for data from inhibitory cells, so that all cells sample from the same cell class for 95% of their profile (all sampled across cells without replacement to ensure there are no confounding overlaps). At each level of rarity and subtlety we measure AUROCs across datasets with MetaNeighbor, using the highest performing GO group for this data as a positive control for gene set selection (identified in the previous analysis to be GO:0022857) and a randomly chosen set of 20 genes as a negative control, having established that small gene sets tend to have low performance.

As expected, GO:0022857 performance is higher than the random set of 20 genes at both 1% and 20% incidences (Figure 3B). Importantly, MetaNeighbor performance is nearly unaffected by differences in rarity: GO set performance is equally high when excitatory neurons make up 1% or 20% of all cells in each dataset, with n as low as 1 cell in the tested data. This is possible because within-dataset labeling is not exploited for training, so rarity is largely irrelevant for scoring. Comparison across multiple datasets in training makes even rare cell types learnable. Of interest is the robustness of MetaNeighbor to transcriptional subtlety. Of course, increasing subtlety leads to worse performance at both incidences, and falls to chance levels at subtletlies >99% (AUROC=0.5). However, even at almost 90% subtlety MetaNeighbor correctly identifies excitatory neurons with a mean AUROC of 0.71. Since this subtlety is relative to the transcriptional variability that exists between inhibitory and excitatory cells, it is quite extreme. Consistent with our previous results comparing performance across all GO functions, this suggests that there are marked and widespread differences in excitatory and inhibitory neuron gene expression, such that even sampling a small fraction of genes (<10%) allows for identification of these two classes. In sum, these results provide strong evidence that MetaNeighbor is robust to differences in rarity, and gives guidance for the interpretation of AUROC scores in light of this factor, suggesting the subtlety of cell identity relative to the outside control.

**Figure 3.**
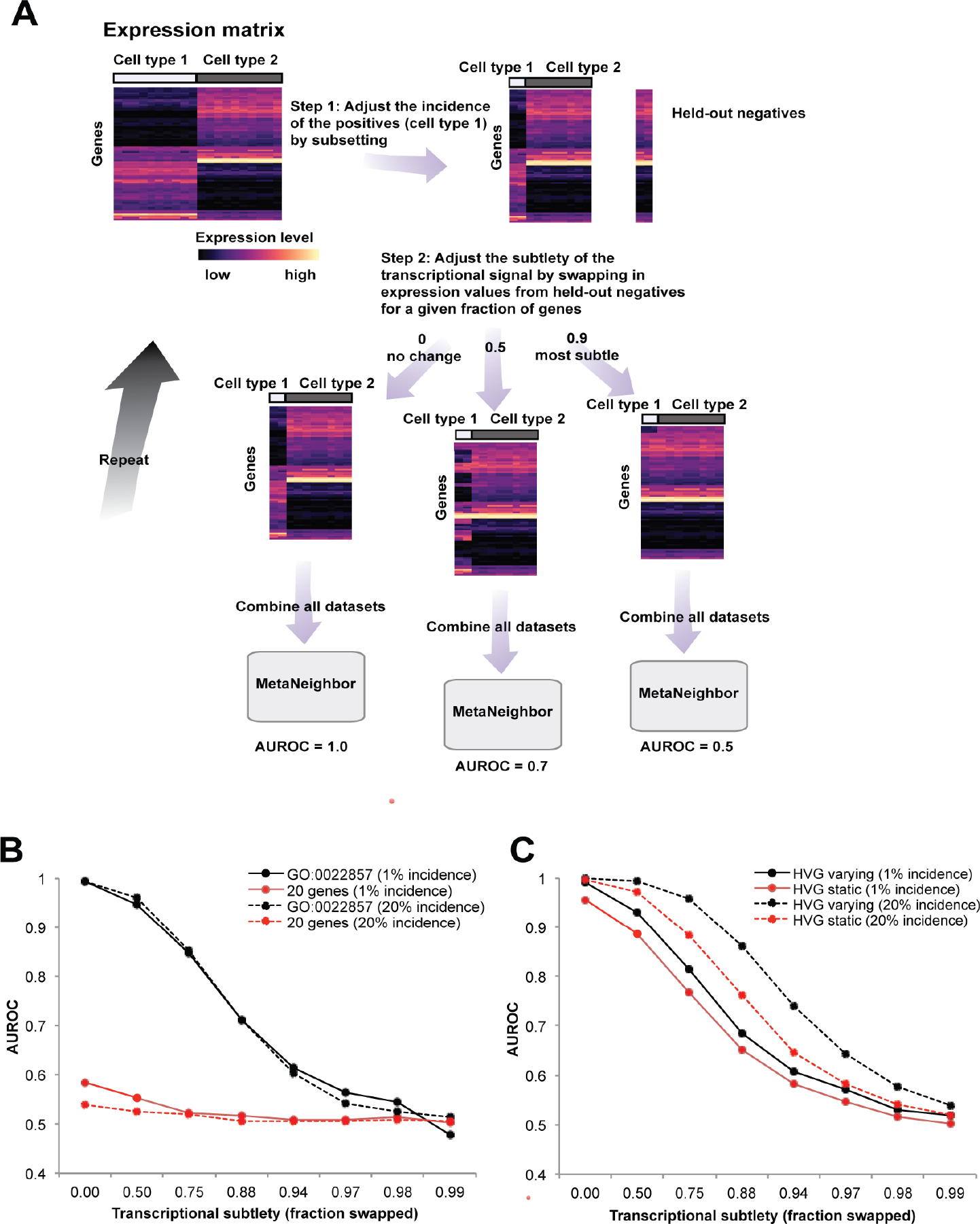
Empirical modeling demonstrates that MetaNeighbor readily identifies rare and transcriptionally subtle cell types. **A** – Schematic of the empirical model. For simplicity only a single dataset is depicted. (Top left) – In this dataset, we begin with an expression matrix containing gene expression levels for two cell types comprising ten cells each. Here we will be assessing the replicability of cell type 1 (‘positives’) relative to cell type 2 (‘negatives’). (Top right) We first adjust cell rarity by randomly sampling subsets of the original expression matrix. In the schematic, incidence is set to 20% (2 positives, 8 negatives). In addition, we partition two negatives from the original data for later use. (Middle) Next, we adjust transcriptional subtlety by randomly sampling genes from a given fraction of the transcriptome. Gene expression in the positives will be replaced with data from the unused negatives, creating a modeled cell type varying from the negative class only in a subset of its genes. (Bottom) All datasets are combined and MetaNeighbor is run to assess the replicability of the positives at each level of rarity and subtlety. **B** – MetaNeighbor results for empirical modeling of excitatory neuron rarity and subtlety, repeated 100 times. Mean performance for the top GO group is in black, performance for 20 randomly chosen genes is shown in red; dashed lines indicate 20% rarity, solid lines show 1% rarity. MetaNeighbor is robust to differences in cell rarity, and can reliably distinguish between types even when they are very similar (AUROC>0.7 at >88% subtlety). **C** – MetaNeighbor results for empirical modeling of excitatory neuron rarity and subtlety using highly variable genes (HVGs), repeated 100 times. Performance for the HVG varying set is shown in black, performance for the HVG static is shown in red; dashed lines indicate 20% rarity, solid lines show 1% rarity. HVGs allow for robust identification of positives even when cells are rare or differences are subtle.

### Empirical modeling to evaluate gene set selection

In the previous section we demonstrated that the highest performing GO group for the excitatory vs. inhibitory comparison is robust to variation in either incidence or transcriptional subtlety, still permitting high-performing identification of these two classes when cells are rare or only subtly distinguishable. Determining this gene set requires known concordance of cell types across datasets. When concordance is unknown, for example when cell type labeling is idiosyncratic, it is necessary to have a strategy to identify informative gene sets *ab initio*. Expert knowledge of informative marker genes is one possibility, though this approach may not be extensible to newly described cell subtypes and suffers from potential ascertainment bias. As a more general alternative, the selection of highly variable genes (HVG) is commonly used in single cell analysis prior to dimension reduction and clustering^4, 7, 33-35^, as it is thought that differentially expressed genes or marker genes should be preferentially variable, and potentially less subject to joint low-level noise. This is in line with our previous observation that gene sets containing highly variable genes are high performing. Indeed, when we select a set of HVG (detailed in Methods) we can almost perfectly identify excitatory neurons compared to inhibitory neurons across datasets (AUROC=0.99) which is equivalent to the highest performing GO group, but without any prior knowledge.

In parallel to our previous analyses, we assessed the robustness of HVG selection at different levels of rarity and subtlety, using either HVG picked from the original dataset that includes all cells (HVG static), or HVG re-calculated based on the precise subset of data included in each run of the empirical model (HVG varying) (Figure 3C). Here, we see that our HVG selection strategy performs equally to or better than the highest performing GO functional gene set for both rare cell types (1%-20% of total), as well as for subtle cell types (differing from out-group by <10%). Interestingly, the HVG heuristic is even responsive to the precise data sampling, yielding modestly improved performance when it is selected based on the precise data generated by the empirical model. It is, perhaps, unsurprising, that the heuristic which many teams of researchers have converged on is a profoundly useful one, but its elegance and robustness are not only valuable but important to understand as a likely baseline upon which more complicated approaches will rest.

These results provide evidence that MetaNeighbor can readily identify cells of the same type across datasets, without relying on specific knowledge of marker genes, even when cells are rare (1% total) or only subtly different from other cells in the out-group against which they are being compared. Importantly, these results also provide guidelines for interpreting AUROCs at cell incidences >=1% in terms of their implications for the promiscuity of cell identity across the transcriptome.

### Investigating cortical interneuron subtypes using MetaNeighbor

Cortical inhibitory interneurons have diverse characteristics based on their morphology, connectivity, electrophysiology and developmental origins, and it has been an ongoing goal to define cell subtypes based on these properties^18^. In a related paper^40^, we describe the transcriptional profiles of GABAergic interneuron types which were targeted using a combinatorial strategy including intersectional marker gene expression, cell lineage, laminar distribution and birth timing, and have been extensively phenotyped both electrophysiologically and morphologically ^42^. Previously, two studies were published in which new interneuron subtypes were defined based on scRNA-seq transcriptional profiles^11, 20^. Because of differences in experimental design and analytic choices, the two studies found different numbers of subtypes (16 in one and 23 in the other). The authors of the later paper compared their outcomes by looking at the expression of a handful of marker genes, which yielded mixed results: a small number of cell types seemed to have a direct match but for others the results were more conflicting, with multiple types matching to one another, and others having no match at all. Here we aimed to more quantitatively assess the similarity of their results, and compare them with our own data which derives from phenotypically characterized sub-populations; i.e., not from unsupervised expression clustering (see Supplementary Table 2 for sample information).

To examine how the previously identified interneuron subtypes are represented across the three studies, we tested the similarity of each pair of subtypes across datasets using HVGs. This was done by alternately considering each subtype as the positive training set, and each other subtype as the test set, answering questions of the class, e.g., “How well does the Zeisel_Int1 HVG expression predict the identity of the Tasic_Smad3 subtype relative to all interneurons in the Tasic data? How well does Tasic_Smad3 HVG expression predict Zeisel_Int1 identity relative to all other interneurons in the Zeisel data?”. Each subtype ranges in incidence from 1-24% of the total number of cells within its own dataset, well within the range of the sensitivity of MetaNeighbor as established above. For each genetically-targeted interneuron type profiled by Paul *et al*., we find a reciprocal best match in the pre-existing data: Paul Sst-Nos1/Tasic Sst-Chodl (AUROC=1), Paul ChC/Tasic Pvalb-Cpne5 (AUROC=0.99), Paul Sst-CR/Tasic Sst-Cbln4 (AUROC=0.98), Paul Pv/Tasic Pvalb-Wt1 (AUROC=0.96), Paul Vip-CR/Tasic Vip-Chat (AUROC=0.96), Paul Vip-Cck/Tasic Vip-Sncg (AUROC=0.95) (Figure 4, all scores in Supplementary Table 3). In addition, expanding our criteria to include all reciprocal best matches, and those with AUROC scores >=0.95, we find additional matches for the Paul subtypes, as well as correspondence among five subtypes that were assessed only in the Tasic and Zeisel data: Tasic Smad3/Zeisel Int14 (AUROC=0.97), Tasic Sncg/Zeisel Int6 (AUROC=0.95), Tasic Ndnf-Car4/Zeisel Int15 (AUROC=0.95), Tasic Igtp/Zeisel Int13 (AUROC=0.94) and Tasic Ndnf-Cxcl14/Zeisel Int12 (AUROC=0.91). Overall we identified 11 subtypes representing 24/45 (53%) types (Figure 4A), with total n for each subtype ranging from 25-189 out of 1583 interneurons across all datasets (1.5-11%). Our corresponding subtypes also confirm the marker gene analysis performed by Tasic *et al*. (Supplementary Table 3), without requiring manual gene curation. Because we quantify the similarity among types we can prioritize matches, and use these as input to MetaNeighbor for further evaluation.

**Figure 4.**
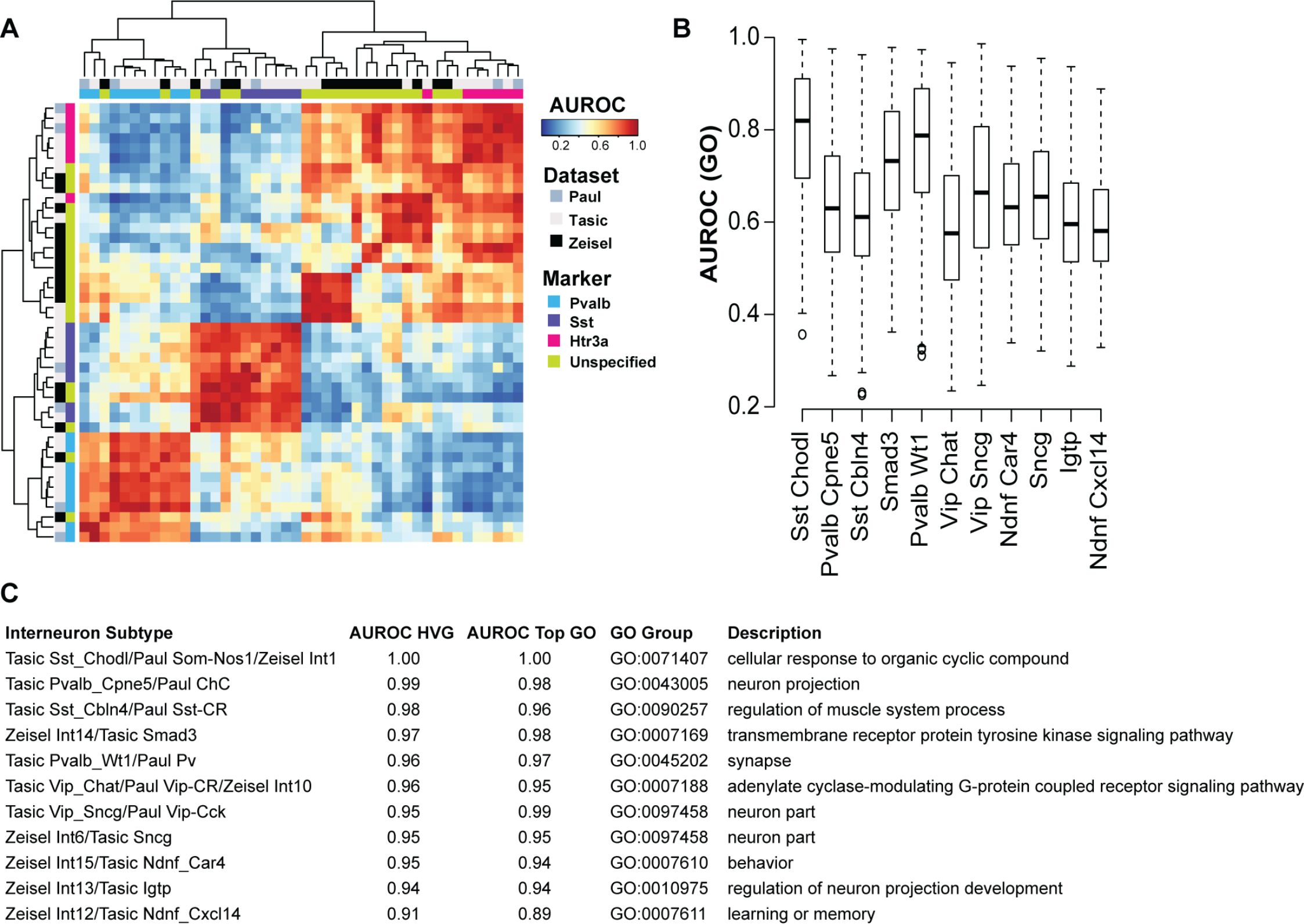
Cross-dataset analysis of interneuron diversity. **A** – Heatmap of AUROC scores between interneuron subtypes based on the highly variable gene set (HVG). Dendrograms were generated by hierarchical clustering of Euclidean distances using average linkage. Row and column colors indicate data origin and marker expression. Clustering of AUROC score profiles recapitulates known cell type structure, with major branches representing the Pv, Sst and Htr3a lineages. **B** – Boxplots of GO performance (3888 sets) for each putatively replicated subtype, ordered by their AUROC score from the highly variable gene set. Subtypes are labeled with the names from Tasic *et al*. A positive relationship is observed between AUROC scores from the highly variable set and the average AUROC score for each subtype. **C** – The table shows the top GO terms for each putatively replicated subtype alongside scores from HVGs. HVGs perform comparably or better than the top ranking GO group for 8/11 subtypes.

**Figure 5.**
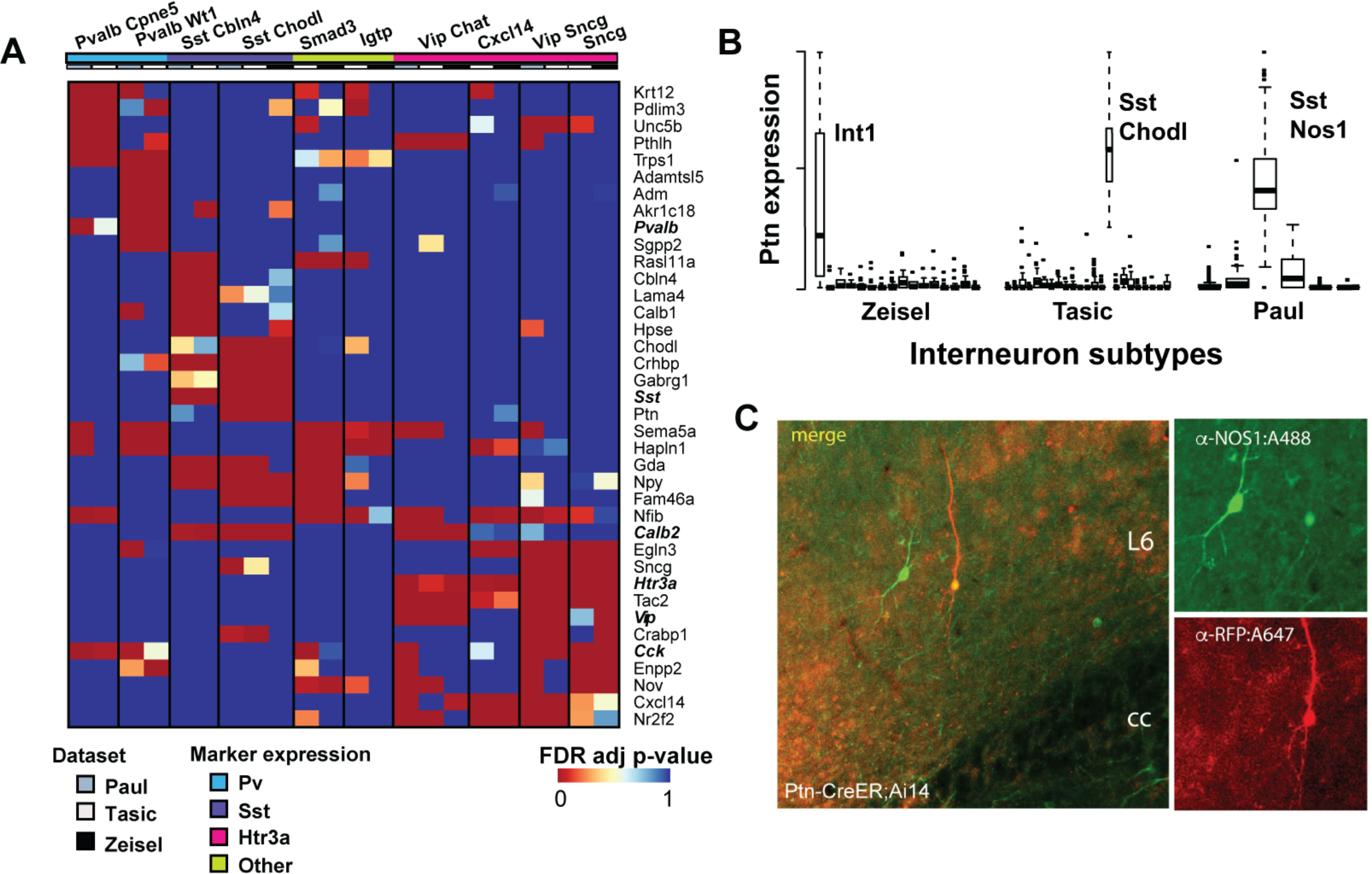
Replicated subtypes show consistent differential expression. **A** – (Top) Heatmap of FDR adjusted p-values of top differentially expressed genes among replicated interneuron subtypes (NB only ten subtypes are shown as no differentially expressed genes were found for the Ndnf Car4 subtype). Subtype names are listed at the top of the columns and are labeled as in Tasic *et al*. Many genes are commonly differentially expressed among multiple subtypes, but combinatorial patterns distinguish them. **B** – Standardized Ptn expression is plotted across the three experiments, where each box represents an interneuron subtype. High, but variable expression is observed across the three Sst Chodl types. **C** – Confocal images of co-immunostaining for Ptn-CreER;Ai14 with RFP and NOS1 antibodies in adult mouse cortex. Ptn-CreER;Ai14 expression was induced with low-dose tamoxifen postnatally. Clear co-labeling is observed in a deep layer (L6) long projecting neuron.

To assess cell identification more broadly, we ran MetaNeighbor with these new across-dataset subtype labels, measuring predictive validity across all gene sets in GO (Figure 4B). The distribution of AUROC scores varied across subtypes but we found that the score from the high variability gene set was representative of overall trends, with high performing groups showing higher mean AUROC scores over many gene sets. Both the high mean AUROCs across all putative replicate subtypes, and the similarity of maximum performance suggest that distinctive gene co-expression can be observed in each subtype (max AUROC=0.92 ± 0.04). As with previous tasks, we found little difference in average AUROCs using functional gene sets compared to random sets (mean AUROC_Random_=0.67 ± 0.06, mean AUROC_GO_=0.68 ± 0.1). Top performing GO groups for each of the 11 replicate interneuron subtypes were primarily related to neuronal function, which is expected due to the large size of these gene sets and their likelihood of expression and variation in these cells (Figure 4C).

These results suggest that highly variable gene sets can be used alongside pairwise testing and training as a heuristic to identify replicable subtypes for further evaluation. Indeed, while outside the scope of our primary analysis, we have found that re-analysis of tens of thousands of cells from mouse cortical and hippocampal pyramidal neurons^11, 12, 20^, retina^6, 7^ and human pancreas^2, 3, 5, 43, 44^ provide strong evidence for the broad applicability of this approach (detailed in the Supplementary Note).

### Identifying subtype specific genes

ScRNA-seq experiments often seek to define marker genes for novel subtypes. Though ideally marker genes are perfectly discriminative with respect to all cells, in practice marker genes are often contextual and defined relative to a particular out-group. Typically, only a very small number of genes are reported in single cell papers due to the complexity of discussing dozens of cell types as well as the potential technical confounds which would limit the expected replicability of any attempt at a more comprehensive list^5, 7, 11, 20^. Here we aimed to identify possible marker genes that would allow discrimination among interneuron subtypes. For each of our identified replicate subtypes we generated a ranked list of possible marker genes by performing one-tailed, non-parametric differential expression analysis within each study for all subtypes (e.g., Int1 vs. all other interneurons in the Zeisel study, Int2 vs. all interneurons, etc.) and combining p-values for replicated types using Fisher’s method (Supplementary Table 4). While data-merging is of potential value in identifying weakly variable genes through improved power, assessing labs independently (“data slicing”) is imperative to identify the most robustly replicable features which will generalize to new labs without additional modeling. Figure 4A shows the FDR adjusted p-values for the top candidates based on fold change for the ten replicated interneuron subtypes with overlapping differential expression patterns. The majority of these genes have previously been characterized as having some degree of subtype-specific expression, for example we readily identify genes that were used for the Cre-driver lines in the Tasic and Paul studies (*Sst, Pvalb, Vip, Cck, Htr3a*), as well as all markers previously reported to intersect between the Tasic and Zeisel data (Supplementary Table 4). Even though we filtered for genes with high fold changes, we see that many genes are differentially expressed in more than one subtype. Notably, considerable overlap can be observed among the *Htr3a-*expressing types. For example, the Vip Sncg subtype (Tasic Vip Sncg/Paul Vip Cck) is only subtly different from the Sncg subtype (Tasic Sncg/Zeisel Int6) across this subset of genes, with the Sncg cells lacking differential expression of *Cxcl14* and *Nr2f2*.

We also identify some novel candidates, including *Ptn*, or pleiotrophin, which is significantly more expressed in the three Sst and *Nos1*-expressing subtypes than in the others (Figure 4B). It is thus expected to be discriminative of these neurons compared to other interneuron types. We validated *Ptn* expression with genetic targeting^40^, and we show clear expression in neurons that stain positively for NOS1 and have morphological features characteristic of long projecting interneurons (Figure 4C). *Ptn* is a growth factor, and we suggest that its expression may be required for maintaining the long-range axonal connections that characterize these cells. These cells are well described by current markers, however this approach is likely to be of particular value for novel subtypes that lack markers, allowing researchers to prioritize genes for follow-up by assessing robustness across multiple data sources.

## Discussion

Single-cell transcriptomics promises to have a revolutionary impact by enabling comprehensive sampling of cellular heterogeneity; nowhere is this variability more profound than within the brain, making it a particular focus of both single-cell transcriptomics and our own analysis into its replicability. The substantial history of transcriptomic analysis and meta-analysis gives us guidance about bottlenecks that will be critical to consider in order to characterize cellular heterogeneity. The most prominent of these is laboratory-specific bias, likely deriving from the adherence to a strict set of internal standards, which may filter for some classes of biological signal (e.g., poly-A selection) or induce purely technical grouping (e.g., by sequencing depth). Because of this, it is imperative to be able to compare data across studies and determine some form of consensus. Indeed, while this work was under review, five manuscripts became available that tackle different aspects of this problem, including robust low-dimensional representation and the use of reference data for cell classification^45, 46^, batch correction using nearest neighbors^22^ and data fusion via manifold alignment^23, 47^. Our paper is unique in its aim and ability to quantify the degree of replicability observable within single cell RNA-seq data, making use of interpretable methods and concrete performance metrics. In this work, we have provided a formal means of determining replicable cell identity by treating it as a quantitative prediction task. The essential premise of our method is that if a cell type has a distinct transcriptional profile within a dataset, then an algorithm trained from that data set will correctly identify the same type within an independent data set.

The currently available data allowed us to draw a number of conclusions. We validated the identity of eleven interneuron subtypes, and described replicate transcriptional profiles to prioritize possible marker genes, including *Ptn*, a growth factor that is preferentially expressed in Sst Chodl cells. One major surprise of our analysis is the degree of replicability in the current data. AUROC scores are exceptionally high, particularly when considered in the context of the well-described technical confounds of single-cell data. We suspect this reflects the fundamental nature of the biological problem we are facing: cell types can be identified by their transcriptional profiles, and the biological clarity of the problem overcomes technical variation. Echoing earlier work on cancer subtyping^30^, we caution that orthogonal data will be required to more firmly establish the biological basis of cell identity; the current estimates must be regarded as optimistic since most clusters are defined from gene expression to begin with. However, the clarity of cell identity is further suggested by our result that cell identity has promiscuous effects within transcriptional data. While in-depth investigation of the most salient gene functions is required to characterize cell types, to simply identify cell types is relatively straightforward. This is necessarily a major factor in the apparent successes of unsupervised methods in determining novel cell types and suggests that cell type identity is clearly defined by transcriptional profiles, regardless of cell selection protocols, library preparation techniques or fine-tuning of clustering algorithms.

Our empirical modeling suggests that this clear signal will permit cell types to be identified down to even greater specificity, but not indefinitely, and some areas of concern within even the present data are worth highlighting. In this work we opted to use the subtype or cluster labels provided by the original authors, in essence to characterize both the underlying data as well as current analytic practices. However, this has limitations where studies cluster to different levels of specificity. This reflects quite real ambiguity about the degree of specificity associated with the term “cell type”. For example, nearly all Pvalb subtypes from the Tasic dataset and the Zeisel Int3 type have AUROC scores >0.9 for the Paul Pv type, as can be seen in the bottom left corner of the heatmap in Figure 4A (Tasic Pvalb_Obox3=0.95, Zeisel Int3 = 0.94, Tasic Pvalb_Tacr3 = 0.94, Tasic Pvalb_Rspo2 = 0.92), suggesting that these may form one larger or more general Parvalbumin-positive type. It is here that the concrete meaning of AUROCs helps. While reciprocal top-hits and AUROCs>0.95 reflect extreme confidence in a highly concordant cell type, more moderate scores are still meaningful. In most domains of biological study, AUROCs>0.9 are extraordinarily high (e.g., ^48, 49^), and we suggest that any such pairing is worthy of discussion and likely reflects real overlaps without indicating replicability. Moving past this point and distinguishing between only subtly different types will be difficult for any analysis, and their discovery will require consideration of appropriate controls and comparisons (e.g., sub-clustering or subset comparisons). The notion of experimental control is built into our scoring method (AUROCs), which by definition is comparing positive and negative cases across the data. As in all classification tasks, choice of an unreasonable out-group or control will generate misleading results, and the closest outgroup is usually the most appropriate. Within our current framework we suggest that a hierarchical approach, moving from broad to subtle categories, will provide a comprehensive, multi-scale view of cell type replicability. We note that our implementation is both robust and fast, but further development of MetaNeighbor and its basic framework may yield improvements (e.g., optimization of feature selection, multi-kernel approaches for cell similarity network estimation, more sophisticated machine learning algorithms).

A key bottleneck, however, is the availability of the data itself. While many groups make their data available in some format, without field-wide standards this data is necessarily more difficult to wrangle than it need be. A common issue is the absence of inferred cell type labels. While it will likely take time and concerted effort for naming conventions to be established, it is crucial that authors make cell labels publicly available in easy-to-access flat text files along with the final parsed expression data matrix to which those labels were applied (or derived). Our wish list for study metadata would also include standardized reporting of cell viability estimates, cell capture method, library preparation method and batch identifiers, alongside biological covariates such as age, sex and strain. More comprehensive reporting would allow for deeper evaluation of technical and biological factors that influence single cell expression results. As the project of assembling a comprehensive human cell atlas gets underway^50^, we hope that participants continue to learn lessons from MAQC and other large consortia projects, making results quickly and readily available to the public, and recognizing the value of heterogeneity in experimental and computational approaches as an assay into biologically robust results with independent and replicable evidence.

## Online Methods

### Public expression data

Data analysis was performed in R using custom scripts^51^. Processed expression data tables were downloaded from GEO directly, then subset to genes appearing on both Affymetrix GeneChip Mouse Gene 2.0 ST array (902119) and the UCSC known gene list to generate a merged matrix containing all samples from each experiment. The mean value was taken for all genes with more than one expression value assigned. Where no gene name match could be found, a value of 0 was input. We considered only samples that were explicitly labeled as single cells, and removed cells that expressed fewer than 1000 genes with expression >0. Cell type labels were manually curated using sample labels and metadata from GEO (see Tables S1 and S2). Merged data and metadata are linked through our Github page.

### Gene sets

Gene annotations were obtained from the GO Consortium ‘goslim_generic’ (August 2015). These were filtered for terms appearing in the GO Consortium mouse annotations ‘gene_association.mgi.gz’ (December 2014) and for gene sets with between 20-1000 genes, leaving 106 GO groups with 9221 associated genes. Random gene sets were generated by randomly choosing genes with the same set size distribution as GO slim. Gene sets based on coefficient of variation were generated by measuring the coefficient of variation for each gene within each dataset, ranking these lists, then taking the average across datasets. The average was then binned into centiles. Sets of highly variable genes were generated by binning data from each dataset into deciles based on expression level, then making lists of the top 25% of the most variable genes for each decile, excluding the most highly expressed bin. The highly variable gene set was then defined as the intersect of the highly variable gene lists across the relevant datasets. Although this did not occur within our analysis, the use of the intersect is likely to be too stringent as the number of datasets for comparison increases. In this case, a majority rule on the highly variable set across datasets appears to be a practicable strategy.

Further commentary regarding high variable gene set selection may be found in the Supplementary Note.

### MetaNeighbor

All scripts, sample data and detailed directions to run MetaNeighbor in R can be found on our Github page ^51^.

The input to MetaNeighbor is a set of genes, a data matrix and two sets of labels: one set for labeling each experiment, and one set for labeling the cell types of interest. For each gene set, the method generates a cell-cell similarity network by measuring the Spearman correlation between all cells across the genes within the set, then ranking and standardizing the network so that all values lie between 0 and 1. The use of rank correlations means that the method is robust to any rank-preserving normalization (i.e., log2, TPM, RPKM). Ranking and standardizing the networks ensures that distributions remain uniform across gene sets, and diminishes the role outlier similarities can play since values are constrained. In previous work we have demonstrated that networks constructed in this way are both robust and highly effective for capturing gene co-expression as evaluated by a variety of machine learning methods^52^.

The node degree of each cell is defined as the sum of the weights of all edges connected to it (i.e., the sum of the standardized correlation coefficients between each cell and all others), and this is used as the null predictor in the neighbor voting algorithm to standardize for a cell’s ‘hub-ness’: cells that are generically linked to many cells are preferentially down-weighted, whereas those with fewer connections are less penalized. For each cell type assessment, the neighbor voting predictor produces a weighted matrix of predicted labels by performing matrix multiplication between the network and the binary vector (0,1) indicating cell type membership, then dividing each element by the null predictor (i.e., node degree). In other words, each cell is given a score equal to the fraction of its neighbors, including itself, which are part of a given cell type ^53^. A difference from KNN is that all cells are neighbors to one another, just to varying degrees (defined by the weighted cell-cell similarity network). For cross-validation, we permute through all possible combinations of leave-one-dataset-out cross-validation, sequentially hiding each experiment’s cell labels in turn, and then reporting how well we can recover cells of the same type as the mean area under the receiver operator characteristic curve (AUROC) across all folds. A key difference from conventional cross-validation is that there is no labeled data within the dataset for which predictions are being made. Labeled data comes only from external datasets, ensuring predictions are driven by signals that are replicable across data sources. To improve speed, AUROCs are calculated analytically, where the AUROC for each cell type *j*, is calculated based on the sum of the ranks of the scores for each cell *i* (*Ranks_i_*), belonging to that cell type, ranked out of all cells within the dataset. This can be expressed as follows:

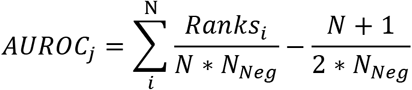

where N is the number of true positives (cells of type *j*), and N_Neg_ is the number of true negatives (cells not of type *j*). Thus, the AUROC calculates the probability that the classifier correctly predicts that a cell of type *j* outranks a cell not of type *j* within the test data set based on similarity to the labeled data in the training data set(s). Note that for experiments with only one cell type this cannot be computed as there are no true negatives. AUROCs are reported as averages across all folds of cross-validation for each gene set (excluding NAs from experiments with no negatives), and the distribution across gene sets is plotted.

### Empirical model of cell type rarity and subtlety

To test the impact of cell type rarity and transcriptional subtlely on MetaNeighbor performance, we repeated the excitatory vs. inhibitory cell discrimination task using the Tasic, Zeisel and Habib datasets which contained >100 cells per cell type, allowing us to assess cell incidences as low as 1%. The essence of the model is to construct a genes by cells matrix in which the biclustering problem to identify cell types from their variation in expression would be increasingly challenging, with both a smaller and smaller fraction of cells (rarity) within the minority class and a smaller and smaller fraction of transcripts distinguishing those cells (subtlety). We model this variability in transcriptional subtlety by sampling different fractions of the transcriptome from the minority class; so, for example, a dataset could be generated in which only 1% of cells have only 10% of their gene expression values sampled from the minority class with the remainder sampled from the majority class. Each minority class cell’s expression vector would thus be the discrete combination of two real cells, one excitatory and one inhibitory. In all cases, real expression values are used with strict partitioning, e.g., sampling without replacement from expression vectors defining cells. Each analysis for a given value of rarity and subtlety was repeated 100 times and means across random sub-samplings of genes and cells are plotted in Figure 3.

### Identifying putative replicates

In cases where cell identity was undefined across datasets (i.e., cortical interneuron subtypes) we treated each subtype label as a positive for each other subtype, and assessed similarity using HVGs. For example, Int1 from the Zeisel dataset was used as the positive (training) set, and all other subtypes were considered the test set in turn. Mean AUROCs from both testing and training folds are plotted in the heatmap in Figure 4. Reciprocal best matches across datasets and AUROCs>=0.95 were used to identify putative replicated types for further assessment with our supervised framework (detailed above). New cell type labels encompassing these replicate types (e.g. a combined Sst-Chodl label containing Int1 (Zeisel), Sst Chodl (Tasic) and Sst Nos1 (Paul)) were generated for MetaNeighbor across random and GO sets, and for meta-analysis of differential expression. While only reciprocal top-hits across laboratories were used to define putative replicate cell types, conventional cross-validation within laboratories was performed to fill in AUROC scores across labels contained within each lab.

### Differential expression

For each cell type within a dataset (defined by the authors’ original labeling), differential gene expression was calculated using a one-sided Wilcoxon rank-sum test, comparing gene expression within a given cell type to all other cells within the dataset (e.g., Zeisel_Int1 vs all other Zeisel interneurons). Meta-analytic p-values were calculated for each putative replicated type using Fisher’s method^54^ then a multiple hypothesis test correction was performed with the Benjamini-Hochberg method^55^. Top differentially expressed genes were those with an adjusted meta-analytic p-value <0.001 and with log2 fold change >2 in each dataset. All differential expression data for putative replicated subtypes can be found in Supplementary Table 4. Details regarding the generation of Ptn-CreER transgenic mice, immunostaining and imaging may be found in Paul *et al*. The image in panel 5C was taken at the same time as those presented in Supplementary Figure 6 of that paper.

## Author Contributions

JG conceived the study. JG, MC, and JH designed experiments. MC and JG wrote the manuscript. MC and SB performed computational experiments. AP performed immunostaining. JH supervised wet-lab data collection. All authors read and approved the final manuscript.

## Acknowledgments

MC was supported by NIH F32MH114501. SB and JG were supported by NIH R01MH113005 and a gift from T. and V. Stanley. Z.J.H. was supported by NIH 5R01MH094705-04, R01MH109665-01 and the CSHL Robertson Neuroscience Fund. A.P. was supported by a NARSAD Postdoctoral Fellowship. The authors would like to thank Paul Pavlidis, Bo Li and Jessica Tollkuhn for their thoughtful feedback on earlier drafts of this manuscript. We would also like to thank the dedicated researchers who have made their data publicly available. Our work would not be possible without their valuable contributions.

